# CryoEM single particle reconstruction with a complex-valued particle stack

**DOI:** 10.1101/2022.07.28.501909

**Authors:** Raquel Bromberg, Yirui Guo, Dominika Borek, Zbyszek Otwinowski

**Affiliations:** Department of Biophysics, The University of Texas Southwestern Medical Center, Dallas, TX, USA; Ligo Analytics, Dallas, TX, USA; Department of Biochemistry, The University of Texas Southwestern Medical Center, Dallas, TX, USA

**Keywords:** particle stack, contrast transfer function (CTF), padding, Ewald sphere, aberrations, single particle reconstruction (SPR)

## Abstract

Single particle reconstruction (SPR) in cryoEM is an image processing task with an elaborate hierarchy that starts with a large number of very noisy multi-frame images. Efficient representation of the intermediary image structures is critical for keeping the calculations manageable. One such intermediary structure is called a particle stack and contains cut-out images of particles in square boxes of predefined size. The micrograph that is the source of the boxed images is usually corrected for motion between frames prior to particle stack creation. However, the contrast transfer function (CTF) or its Fourier Transform point spread function (PSF) are not considered at this step. Historically, the particle stack was intended for large particles and for a tighter PSF, which is characteristic of lower resolution data. The field now performs analyses of smaller particles and to higher resolution, and these conditions result in a broader PSF that requires larger padding and slower calculations to integrate information for each particle. Consequently, the approach to handling structures such as the particle stack should be reexamined to optimize data processing.

Here we propose to use as a source image for the particle stack an exit-wave-reconstruction-based image, in which CTF correction is implicitly applied as a real component of the image. The final CTF correction that we later refine and apply has a very narrow PSF, and so cutting out particles from micrographs that were approximately corrected for CTF does not require extended buffering, i.e. the boxes during the analysis only have to be large enough to encompass the particle. The Fourier Transform of an exit-wave reconstruction creates an image that has complex values. This is a complex value image considered in real space, opposed to standard SPR data processing where complex numbers appear only in Fourier space. This extension of the micrograph concept provides multiple advantages because the particle box size can be small while calculations crucial for high resolution reconstruction such as Ewald sphere correction, aberration refinement, and particle-specific defocus refinement can be performed on the small box data.

**Highlights:**

▪ A complex-valued particle stack facilitates flexible and fast data processing
▪ A real-valued, compact particle stack can be used at intermediate steps
▪ Whole micrograph padding is used from the beginning of data analysis

## Introduction

The SPR method in electron microscopy creates a 3D reconstruction of a particle from a number of projection images whose orientation is *a priori* unknown (“random”) (De Rosier and Klug 1968). This technique has evolved toward high resolution reconstruction starting with very noisy data, with a signal-to-noise ratio (SNR) for most contributions from individual particle images being well below one. For this reason, the number of particle images used in reconstruction can be high, with the use of millions of particles becoming a routine approach in solving challenging projects (Naydenova, Muir et al. 2021, Singh, Vanden Broeck et al. 2021). The analysis that is typically applied to these large image sets is iterative and the particle images are processed many times. Finding an appropriate reduced representation for these images is necessary for the efficiency of cryoEM data processing.

One such reduced representation is the particle stack, where particles are cut out from raw micrographs and placed within uniformly sized boxes (Fig. 1). These boxes require a margin to account for uncertainty of particle positions but also to accommodate the point spread function (PSF) of the imaging system (Fig. 1D). The phase contrast method, used in cryoEM SPR, has an unusually large PSF for an imaging system, even approaching the size of the collected image (Glaeser, Hagen et al. 2021). This is because experimentalists acquire cryoEM data far out of focus to increase the image contrast at low resolution, which is critical for particle identification, but enlarges the size of the PSF. The radius of the PSF depends on the resolution limit and is proportional to the product of the defocus and the resolution, expressed in inverse Angstrom:

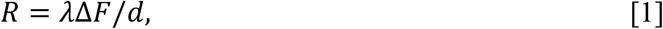

where *R* is the radius of the PSF at a given resolution, λ is the electron wavelength, Δ*F* is the defocus, and *d* is the resolution in Angstrom (Rosenthal and Henderson 2003, Moriya, Adachi et al. 2020, Weis and Hagen 2020).

**Figure 1.**
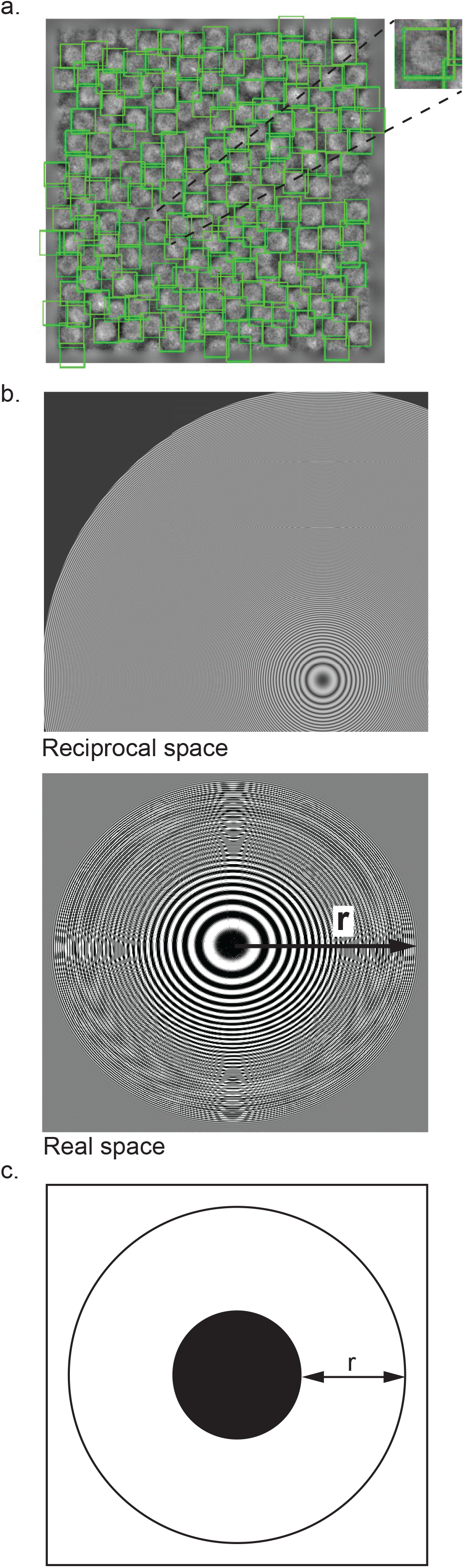
Possible inefficiencies during the creation of the particle stack. (a) Apoferritin micrograph with particles extracted within tight boxes. Even here, the substantial redundancy in areas belonging to multiple boxes can be observed. (b) Top: CTF function for micrograph in (a), in reciprocal space. The number of Thon rings can be high, depending on the focus, voltage and resolution. Bottom: The point spread function (PSF), which is the Fourier Transform of CTF (above). (c) Schematic of how the PSF with radius r creates a particle image on the micrograph that needs to be boxed. This buffer area around the particle (dark filled circle in the schematic, without PSF contribution) can be much larger than the particle projection area. If applied to data from (a), the boxes will overlap grossly and require storage space that is nearly 9× the size of the original micrograph.

The amount of defocus is influenced by the need to identify particles, and so larger values of defocus are used for smaller particles which are more difficult to detect. The impact of the recent “resolution revolution” is that users also now perform reconstructions at higher resolutions. For these reasons, the balance between the size of the PSF and the particle size has changed dramatically. Historically, the size of the PSF was much smaller than the particle size, and so the needed margin around the particle in the box was small. However, now this margin can be much larger than the size of the particle and most of the box image consists of the particle’s surroundings (Fig. 1D), which are needed for CTF deconvolution which is applied to the Fourier Transform of the boxed image (Tegunov, Xue et al. 2021). We call the result of this deconvolution, or some approximation of it, a “CTF corrected” image (Downing and Glaeser 2008, Glaeser, Hagen et al. 2021). The particle stack resulting from these calculations and consisting of all boxed particles can become much larger than the micrographs from which the boxes were cut out of and consequently calculations become expensive. The size of the particle stack becomes even more important when highly parallel graphical processor units (GPUs) are used to speed up calculations because gains from using GPUs can be erased if repetitive reading of data from mass storage is required (Zivanov, Nakane et al. 2018).

From early SPR programs to current ones, a particle stack has been used as an organizing structure stored on disk and facilitating communication between different programs. Recently, Tegunov made the cutout particles an internal, dynamic structure (Tegunov, Xue et al. 2021); this solved the problem of a user having to guess what would be the final resolution needed to calculate the necessary size of the box in the particle stack (Fig. 1D). Our approach goes one step further by applying CTF to the whole micrograph, so cutting out particles does not have to extend significantly beyond the expected particle size. However, cutting particle images out of CTF-corrected images requires modifications in data analysis, in particular if we want to create **a persistent data structure** of CTF-corrected particle images in a stack. We discuss these modifications and how they affect multiple steps of the SPR workflow.

## Results

### The Case of Fixed CTF, Determined Prior to Image Analysis, from the Power Spectrum

In the standard workflow, CTF is determined from the power spectrum (Penczek, Fang et al. 2014) early on and is assumed to be the same both across the entire micrograph and also during the iterative SPR process (Grant, Rohou et al. 2018). Even if the CTF changes later, by applying reference-based refinement (Bromberg, Guo et al. 2020, Zivanov, Nakane et al. 2020), most of the SPR calculations are performed with the initial CTF values. Therefore, using images corrected from the start for CTF is a sensible idea and was considered earlier (Penczek, Fang et al. 2014). However, implementing such a solution creates complications.

The first complication arises from interactions of CTF correction with noise filtering of the image. The CTF correction can be considered a deconvolution of the PSF (Fig. 2). Deconvolutions are unstable with respect to noise, and so the noise filters have to be involved and they are adjusted as our knowledge changes. A typical noise filter applies Wiener weights (Glaeser, Hagen et al. 2021) to structure factors:

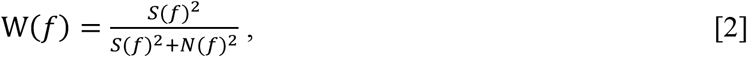

where W is Wiener filter weight as a function of spatial frequency *f*, *S*(*f*) is the current estimate of signal strength including CTF multiplicative contribution, and *N*(*f*) is our estimate of noise. Thus, CTF as a multiplicative normalizer of our measured signal is included in the filter that is needed to perform CTF deconvolution. This leads to an approach where the Wiener filter is applied in two steps, where the multiplication by CTF^2^ is applied early on in data processing (Downing and Glaeser 2008, Glaeser, Hagen et al. 2021), while the remainder of the Wiener filter is applied much later. The advantage is that multiplication by CTF^2^ stabilizes deconvolution, as a division by small values (including zero) is avoided if deconvolution of CTF is combined with multiplication of CTF^2^. The result of this operation is called a CTF-corrected image, which in real space corresponds to convolving the starting image with the PSF. This noise filtering creates the non-intuitive situation where the start of the inverse process is applying the forward part of the data model (Fig. 2).

**Figure 2.**
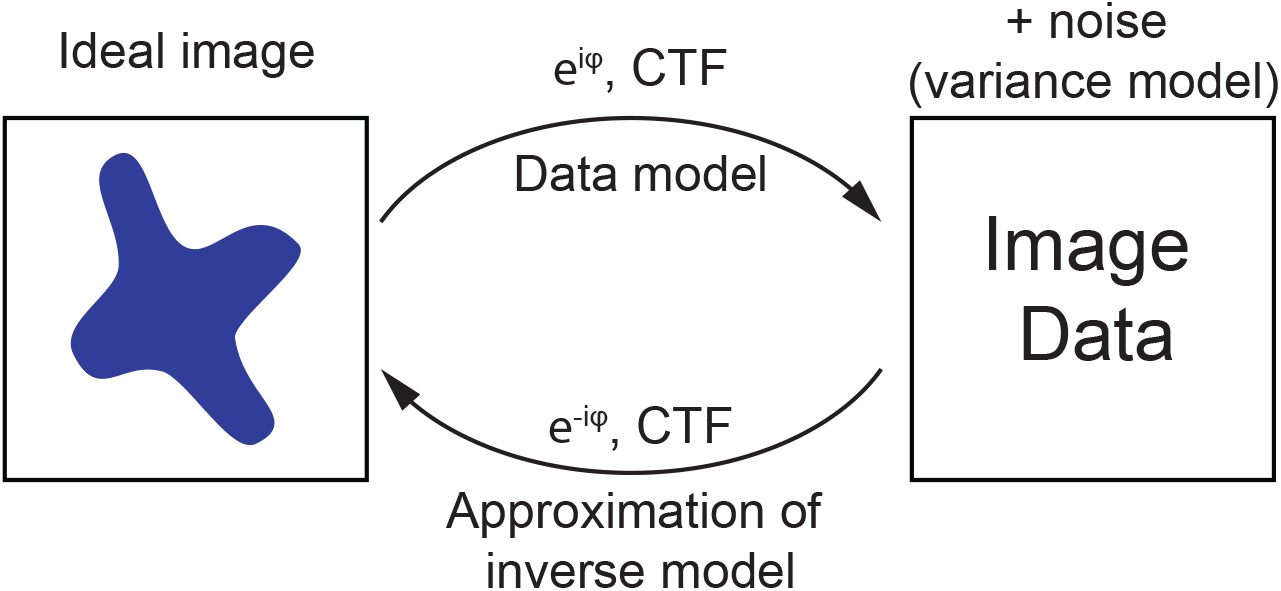
The relationship between the image and the data has a central contribution from wavefront propagation *e^iφ^*, which includes all aberrations and defocus. However, for most data, the thin sample approximation is valid, and then the real part of the wavefront propagation creates the contrast transfer function (CTF). While the inverse model should in principle have CTF^−1^, the high level of noise generates weights which are approximated by CTF^2^, so when multiplied together, they generate a contribution from multiplication by CTF in the inverse model.

The second complication arises from this convolution being performed in reciprocal space and then followed by the Fast Fourier Transform (FFT), which is necessary for speed requirements. The FFT requires periodic boundary conditions which create wraparound effects when performing convolution with extended objects (e.g. PSFs). To avoid the increased noise due to wraparound effects, a buffer needs to be formed around the original image by a process referred to as padding, where the buffer is filled with a background value. Padding has additional consequences that we discuss further below.

### Padding and Binning

When the box around a particle extends outside the micrograph (Fig. 3A), the fixed dimensioning of the box requires filling in the missing data with some type of background estimate. This extension of the original micrograph is an important step in cryoEM workflows, including ours. However, its implementation needs to consider properties of CTF and FFT. CTF correction involves the FFT whose speed is inherently dependent on assuming that the input image has circular (wraparound) symmetry. Convolution with the PSF will extend past the original micrograph boundaries; therefore, to prevent wraparound artifacts, it is necessary to pad the original micrograph with a buffer that is larger than the radius of the PSF. It is convenient to perform padding which is symmetrical between the left and right and between the top and the bottom, but this is not required. What matters is the size of the padding, which is the sum of the left and right and also the sum of the top and the bottom. These sums do not have to be equal if the image is rectangular, but the smaller one should exceed the size of the PSF. This freedom in adjusting padding upwards allows us to choose the size of the padded image size in pixels that optimizes the FFT speed. For GATAN detectors, which are widely used and have image sizes that are inefficient for the FFT in both *x* and *y* directions, padding by the right amount can speed up the FFT calculations by a large factor. For instance, for the K2 detector, padding upwards to 4096 (for *x* and *y*) reduces the number of floating-point operations by ~140×.

**Figure 3.**
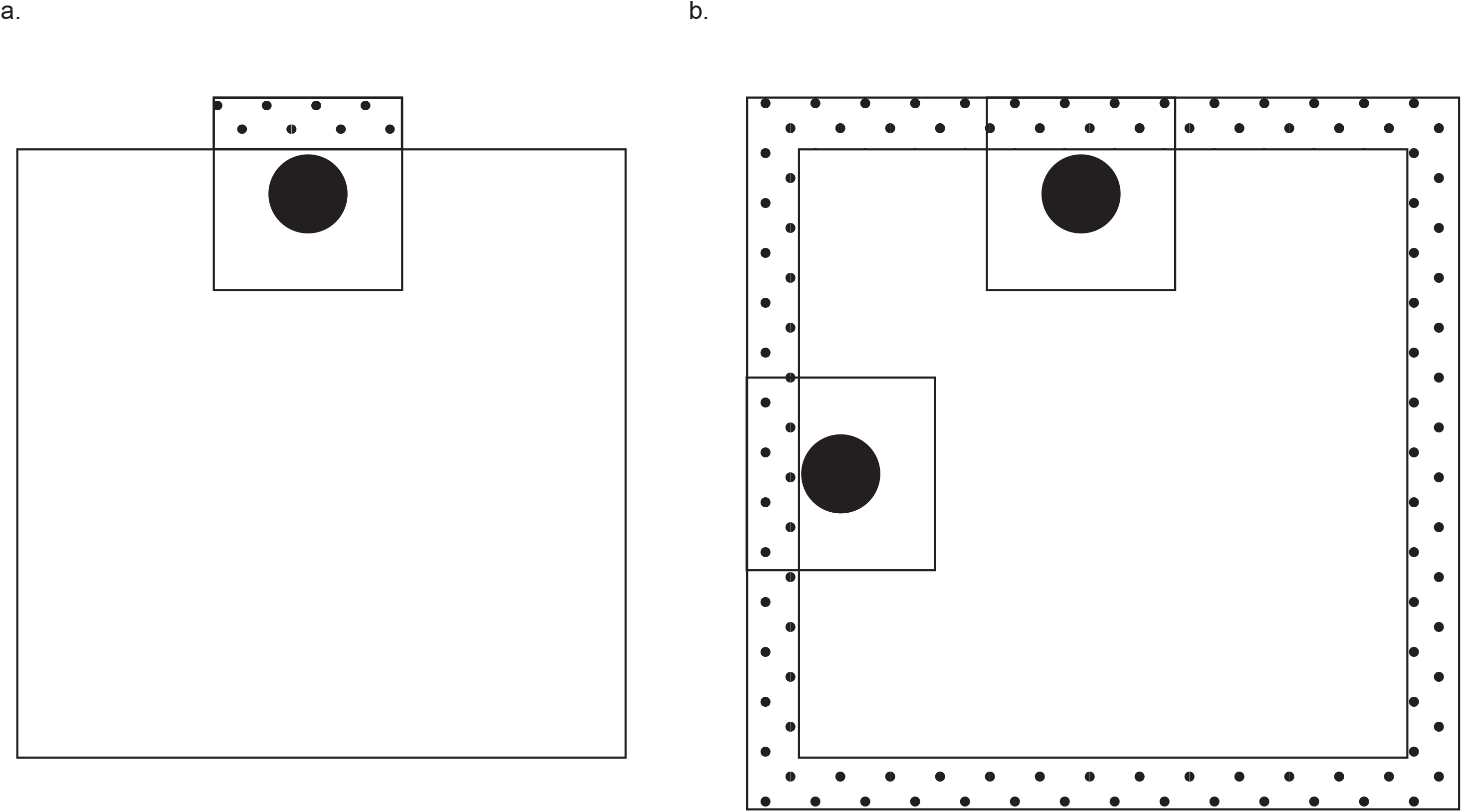
Padding boxes or padding micrographs. Dotted areas represent padding. (a) The traditional workflow has micrographs in which the image data extends from edge to edge (no padding). When a particle box extends outside of the micrograph, the extended area is filled with background values: padding is introduced at the level of the particle box. (b) For micrographs to be CTF-corrected, they need to be padded first. As long as the particles are found within the valid image data, there is no need to introduce padding during particle box extraction. Additionally, the box is likely to be smaller due to CTF correction.

Additional improvements in both speed and accuracy can be obtained by performing this padding on raw movie frames prior to alignment. The speed improvement comes from the faster FFT, described above, although this factor is only true for data from GATAN detectors, as Falcon images factorize well in FFT. The accuracy improvement results from avoiding wraparound effect artifacts when performing motion correction.

Detector quantum efficiency decreases as the spatial frequency of the measured data approaches the detector’s Nyquist frequency, i.e. higher resolution signals are always measured less accurately. This effect is mitigated by increasing the magnification of the microscope, so the highest frequency of interest (inverse of resolution) corresponds to a fraction of the Nyquist frequency. This creates an oversampling condition that is detrimental to data processing efficiency. Therefore, it is more convenient to reduce data resolution by a procedure analogous to binning but performed in Fourier space. This procedure fully preserves a high frequency signal up to the Nyquist frequency of this “binned” data by performing Fourier space truncation (cropping). An additional advantage of Fourier space truncation is that it allows for precise binning without interpolation, by not only integer numbers, but also rational fractions. However, binning by rational fractions for rectangular detectors may produce slightly different binning factors for the *x* and *y* axes due to FFT speed considerations, resulting in rectangular (but almost square) pixels. Fortunately, Zivanov et al. provided a formulation for how to handle rectangular pixels [Eq. 24 in (Zivanov, Nakane et al. 2020)] in downstream calculations, and we use it both for anisotropic magnification and also for different binning factors along the *x* and *y* axes. The 2×2 matrices that describe these linear transformations need to be multiplied to produce the matrix for the unevenly binned data.

### Consequences of our modified padding, binning, and CTF-correction

We introduced padding starting from movie frames to optimize FFT and prevent wraparound effects in motion correction and also subsequently in CTF correction. We then applied CTF correction and Fourier truncation to reduce the resolution of the micrographs, while at the same time optimizing the image size to speed up the FFT. The CTF correction reduced the requirement for the box size of the particles that are cut out from such micrographs, reducing the number of calculations in classification and 3D reconstruction. In our modified workflow, there is no requirement to create a persistent particle stack (Tegunov, Xue et al. 2021), which reduces the I/O load for images with high density of particles (Fig. 1).

### Variable CTF Across a Micrograph and/or Defocus-Within-Particle Correction

A particle has a volume, so the defocus associated with the particle varies within the range defined by the size of the particle. Defocus variation within a reconstructed volume from an individual particle image (DeRosier 2000) is a direct consequence of the core physics of scattering and measurement. We are interested in energy-preserving elastic scattering that is represented by the surface of electron momentum change—the Fourier-space Ewald sphere of energy conservation. Non-flatness of the Ewald sphere means the measured image is not a direct projection but is perturbed by variations in the defocus at high resolution. Accurate handling of this defocus variation, so-called Ewald sphere correction, has two requirements. The first is that in reconstruction, the contribution from a particle image is a curved surface (Ewald spheres, Fig. 4) rather than a plane. The second is that we insert into the reconstruction the image which has been corrected in reciprocal (Fourier) space by a complex function describing the defocus and aberrations. Such a corrected image is closely related to the image that would result from the exit-wave reconstruction process (Wolf, DeRosier et al. 2006, Chen, Schmidt et al. 2021). In macromolecular cryoEM, the Ewald sphere correction matters only for resolutions at which the SNR is well below 1.0. Under such conditions, the contribution to the 3D reconstruction arises from the product of the Fourier Transform of the data (micrograph) and the wavefront function:

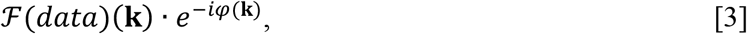

where 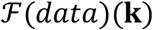 is a Fourier transform of a micrograph at momentum **k** on the Ewald sphere and *φ*(**k**) is the wavefront function. The contribution from the wavefront function is not Hermitian, so the Fourier Transform of this product is a complex value image. This complex value image has a real part that corresponds to the traditional CTF correction performed on a plane, but also has an imaginary component that describes the CTF correction where the cosine function is replaced by a minus sine function. This complex value image is placed on the Ewald sphere, and this step describes the quantum mechanical (QM) wave function resulting from scattering (Fig. 4). The measurement process involves a collapse of the wave function, and the magnitude of the wave (which is what we can measure) involves both the scattered wave function and its complex conjugate. This complex conjugate creates a contribution from a Hermitian copy of the Ewald sphere (Wolf, DeRosier et al. 2006, Russo and Henderson 2018). The Hermitian copy means 3D inversion through the origin, combined with applying the complex conjugate (Fig. 4). Together, the contributions from both spheres are inserted into the 3D reconstruction space. When these contributions are highly correlated, the imaginary component of the image, originating from the sine term, cancels out, and the remaining real component of the wave function generates the CTF modulation of the power spectrum. We propose to separate the description of the scattering and measurement processes in our data structures.

**Figure 4.**
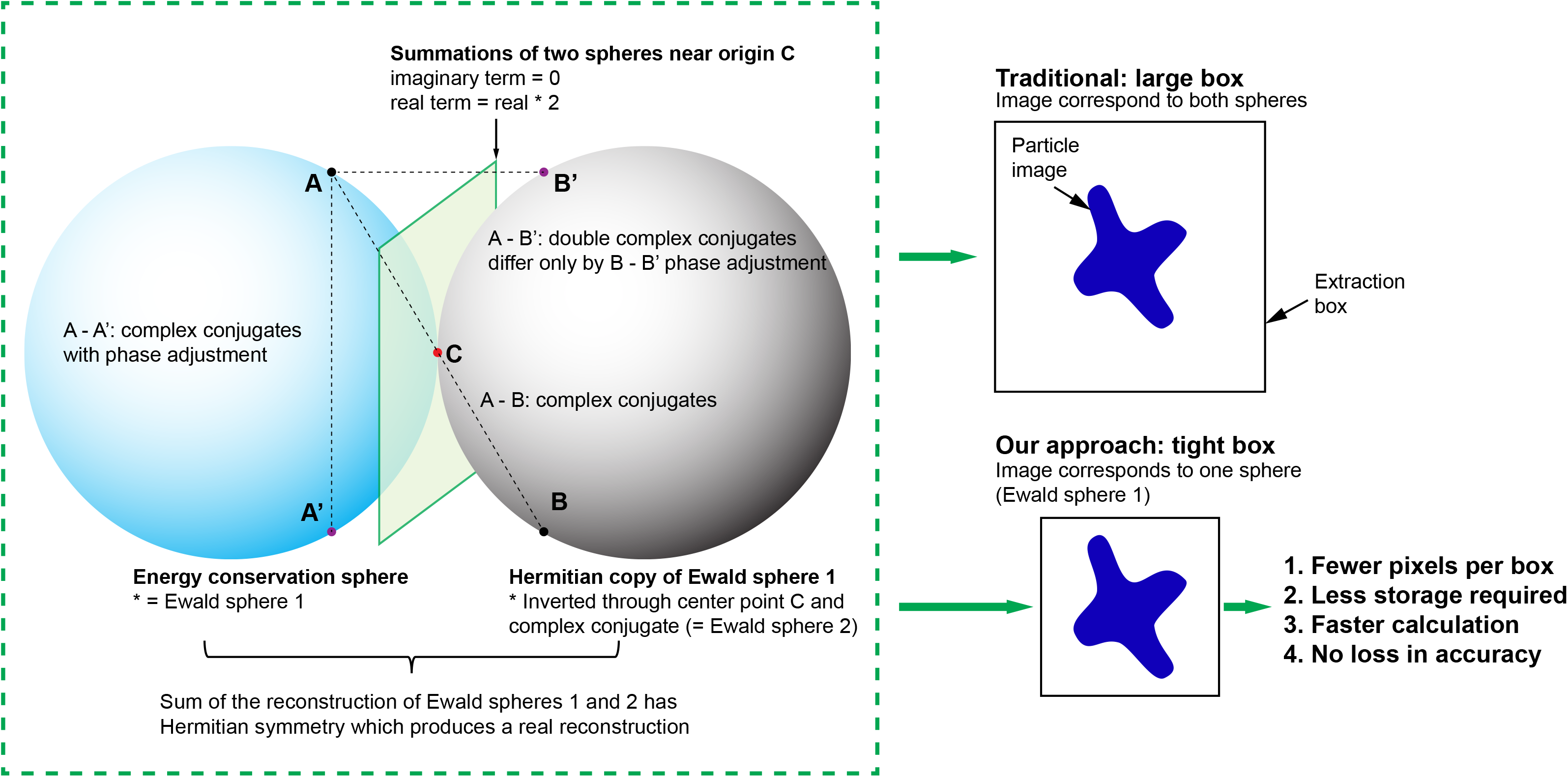
Graphical Abstract: The classical approach in cryoEM SPR merges image phase adjustment with 3D reconstruction, first in a thin sample approximation, and then with corrections for departure from this approximation. In our approach, we start with precise Ewald sphere formulation: we first adjust the phases in the image, possibly via a hierarchical procedure, unlike the classical approach. We consider amplitude, which affects weights, only in the 3D step, where contributions from the Ewald sphere and its Hermite mate are merged. In the classical approach, any change in the phase adjustments requires going back to the starting micrographs, as phase adjustment (i.e. aberration refinement) cannot be stacked. This creates a computational inefficiency.

Splitting the image formation process into wavefront propagation and the QM measurement process is inherent to Ewald sphere correction. While Ewald sphere correction is not needed at low resolution, creating data structures that support this split in reconstruction has advantages at lower resolution as well. The primary data structure corresponds to inversion of the wave propagation applied to the observed image (Fig. 4) and can be used in the final stages of reconstruction to refine the CTF to account for aberrations, including defocus per particle. These refinements introduce small changes to the CTF, but the traditional CTF-corrected image is not suitable as a starting point for CTF adjustments, so standard procedures rely on the uncorrected particle stack. The exit-wave-reconstruction-type data structure we describe here allows for very convenient stacking of CTF correction. These corrections are small, and so have a very narrow PSF, and consequently applying them to boxed images does not require an enlarged buffer area around the particles. In particular, we can apply the component which has the largest PSF first, calculate our complex value image, save the boxed data as complex value images, and apply corrections with a narrow PSF later.

Creating an intermediate structure as a complex image, either at the level of a micrograph or a particle stack, has three benefits. First, we do not have to oversize the particle cutouts (e.g. boxes or circles). Second, we can perform additional CTF corrections on these images, which is particularly important for refinement of individual particle defocus, for correcting defocus introduced by the tilt of the sample, or for other even-order aberrations. Third, we can correct Ewald sphere curvature (variable defocus-within-particle or tomograph) during 3D reconstruction, with minimal complexity.

The penalty we pay for this scheme is that the complex value images are twice as large in storage, when the number of pixels is the same. For this reason, we should create either real-value or complex-value images depending on these defocus-variation considerations and how they apply to specific calculations. Usually, these calculations will have associated specific resolution limits, so these considerations go together. For example, images for the purpose of particle picking can be real, images for the purpose of 2D classification can almost always be real, and images for 3D reconstruction could be real or complex depending on the need for defocus (Ewald sphere) correction, other CTF corrections, particle size and resolution. We note that stacking defocus corrections using these complex value images simplifies and accelerates calculations of variable defocus-corrected images for tilted samples. Such images still allow for calculating further corrections, like aberration refinement and individual particle defocus refinement.

### Wiener-like weights during reconstruction

So far, we have discussed CTF correction only as an approximation to the Wiener filter. However, at some point in the calculations, we would like to apply the proper Wiener filter (Glaeser, Hagen et al. 2021). Here we differentiate cases where particles are (1) averaged, e.g. during 2D class classification, and are (2) not averaged, e.g. when the Wiener filter is applied to a micrograph. For the latter case, the useful resolution limit is low, so the Ewald sphere curvature considerations do not apply. As discussed above, we apply the remainder of the Wiener filter to the CTF correction. Then we calculate the Wiener-filter-corrected image by backwards FFT.

We now discuss the former case. Averaging multiple particle images involves calculating a weighted average, which needs to be done in Fourier space because this is where the weights factorize. Applying the Wiener filter is the last step in this process, both for 2D classification and 3D reconstruction.

The averaging of information from images with different CTFs involves hierarchical, multi-step reasoning, for which obtaining the resolution-dependent SNR estimates is central. The best resolution-dependent SNR estimates are created by comparing two half-reconstructions and calculating the weighted FSC between them (Sindelar and Grigorieff 2012). We start with the reciprocal space accumulation of contributions from weighted images (CTF-corrected) and the sum of weights (the sum of CTF^2^). Summation of the CTF-multiplied images is straightforward, as the PSF size effects were already accounted for. Summation of CTF^2^ involves only positive values and generates a scaling factor for the sum of the CTF-corrected images, which have large relative error. Summation of CTF^2^ is a challenge because in principle, it requires a box that is twice as large as defined by PSF considerations. However, because of the non-stringent requirements for calculating it accurately, we can approximate CTF^2^ with an unbiased estimator which can be used even if it is sub-optimal, because it contributes little to the uncertainty of the results. This allows for multiple approaches to calculating the sum of CTF^2^. What is more accurate but also computationally involved is calculation of the CTF^2^ term in the space that corresponds to the Fourier Transform of an enlarged box which is twice as large as the PSF limit (Fig. 1D) (Tegunov 2021). Subsequently, we take the Fourier Transform of the CTF^2^, filter it in real space so that it does not extend much past the particle radius, and then Fourier Transform it back to generate a filtered CTF^2^ function. Faster, more *ad hoc* methods to suppress CTF^2^ oscillations can also be suitable (Penczek, Fang et al. 2014).

In the case of 3D reconstruction, the remaining step of back projection depends on the details of algorithms for dual-space filtering in different programs. This dual-space refers to Wiener filtering in reciprocal space and particle-size-related filtering in real space; the combined filtering does not have a simple optimal solution. Therefore, various computational simplifications are being used to avoid excessive calculations in more optimal algorithms (Sindelar and Grigorieff 2012). Intertwined with these simplifications is a question of interpolation errors, both in real and reciprocal space (He, Govindaraju et al. 2007, Penczek, Fang et al. 2014, Střelák, Sorzano et al. 2019). How to handle all these tradeoffs is an under-discussed subject, and implementations may hide these decisions. Wiener filtering is compatible with our CTF-corrected images of reduced box size around particles, but with some variable consequences depending on how the Wiener weights propagate through 2D/3D reconstruction programs in actual implementations.

One should note that in the above considerations, the box size impacts only the back projection step rather than the orientation refinement. Particle refinement does not involve averaging, so it needs only weight correction at low resolution, discussed earlier for micrographs. The more computationally involved CTF^2^ term appears only during calculation of the multi-particle average. As particle refinement is the slowest step in the SPR process, the potential PSF-size-driven inefficiency of calculating the CTF^2^ contributions is not very consequential.

## Discussion

Here we described how complex-value images can be used in the 3D reconstruction process in SPR to better organize calculations. The quantum mechanical description of the measurement process imposes a Hermitian symmetry which in our approach is applied only in 3D back projection (Wolf, DeRosier et al. 2006). Our formulation is fully 3-dimensional, where the experimental 2D image is formed on the surface of the Ewald sphere. Using 2D images which, as in the exit-wave reconstruction, have complex values has multiple consequences in terms of speeding up and simplifying calculations. Our data structure (complex value image) represents the QM state prior to the measurement process, and so it has fundamental significance and is not just a computational simplification from using simple complex-number algebra. In addition, if we use the CTF-corrected complex image as an intermediary in calculations, in particular when creating particle boxes, the subsequent calculations can be significantly faster due to substantial reduction in the number of pixels that need to be analyzed, as the boxes have become much smaller.

When creating complex value 2D images from micrographs, only a phase shift factor (Eq. 3) and no amplitude factor is applied. The amplitude factor, known as the CTF, arises from the measurement step description in 3D space, where the real and imaginary parts of the 2D image become correlated and anti-correlated in 3D, respectively. The amount of correlation is a function of sample thickness and resolution. For thin samples, the strong anti-correlation of the imaginary component reduces its 3D contribution to almost zero, while the real component generates well-known (Thon rings) amplitude modulation. The Ewald sphere correction represents the case where both correlation and anti-correlation of the image components approach zero (i.e. sample is no longer considered to be thin). We note that the use of CTF weights in Wiener filtering is a consequence of a thin sample approximation and needs to be adjusted at higher resolution. In our 2D complex image approach, we disentangle phase corrections which can be stacked from the amplitude modulation aspect which comes at the end of the calculations during the 3D step. In terms of getting phases and amplitudes right, getting the phase right is crucial, while amplitude considerations represent only an optimization of the SNR in the image. For this reason, when we separate these two aspects, the phase-adjusted complex value 2D image becomes the pivotal intermediate step in the analysis.

The current workflows of SPR involve several core procedures: movie alignment with resolution dependent correction for signal decay in individual frames, initial CTF analysis based on the power spectrum, image interpretation such as particle picking and artifact identification, and multiparticle analysis that involves averaging, such as 2D classification and 3D reconstruction by back projection where the averaging is performed in reciprocal space. These steps provide progressively better SNR, which at the start in individual movie frames is too low for anything to be recognized, micrographs being borderline interpretable. Subsequently, CTF analysis allows for Wiener filtering which makes micrographs more interpretable, and only averaging makes features at atomic or close to atomic resolution identifiable.

Experimental datasets are now reaching the order of one million particles, and in the averaging steps, involve the complex tasks of classification or 3D reconstruction where computational efficiency is paramount. In addition, there is a tradeoff between the speed of calculations and the quality of the approximations used in the process. This technical aspect is highly significant, as this is the step where the data become interpretable but the calculations can become expensive and/or difficult to manage due to the size of the data and the lack of a clear path through the calculations under frequent conditions such as preferred orientation, multiple conformations, small particle size, etc.

CryoEM SPR involves measurement under an unusually large (for optics) PSF and this has multiple consequences during the reconstruction process, which we discussed in detail above. Another unusual aspect for image processing is using images with very low SNR for most of the resolution range of the result. We have inherent limitations on how much SNR we can achieve when measuring individual macromolecules, with SNR of about 1 extending only to resolutions in the vicinity of 15-20 Å. This is far below what is needed for building atomic models, and to reach the resolutions enabling it, e.g. 3-4 Å, extensive averaging of multiple particles is needed. Filtering out noise is the most important consideration here. The CTF can also be considered a multiplicative normalizer of the data; it defines our signal model as a product of a particle’s Fourier Transform and the CTF function, which can be different for every particle. Correcting for CTF is a form of deconvolution where due to the frequent zeroes of the CTF function, noise regularization is required with a reciprocal space Wiener filter being the first layer of noise suppression. We have elements of a bootstrap problem, as we need a Wiener filter to produce a reconstruction, and we need a reconstruction to produce a signal model. The signal model is not fixed in the process, as it may increase when alignment of particles becomes more accurate due to the reference becoming more informative in consecutive iteration cycles. In addition, our original CTF estimates may be improved by better refinement of aberrations, when the reference becomes available (Bromberg, Guo et al. 2020, Zivanov, Nakane et al. 2020).

In this paper, we have discussed improved approaches for organizing calculations in the presence of these rather typical features of reconstruction. In particular, we described a replacement for the particle stack, a traditional data structure where images of particles were stored as separate items in boxes of some fixed size. The improved approach allows for tighter margins around particles and involves the unusual feature of creating real space complex images. The real part of these complex images has the same meaning as in traditional images. However, storing the imaginary part allows for refining and/or adjusting the CTF using a particle-sized image, without the need to “reprocess” the whole micrograph or a large box. The number of calculations in SPR is mostly driven by the total number of pixels in the particle boxes. Therefore, making tighter boxes significantly increases the computational efficiency of the process.

Adjusting the defocus represents a fundamental aspect of reconstruction, and in principle, defocus is variable even within a single particle. In other words, the images are not plain projections, which is a feature of the simplified presentation of the measurement process. Instead, the Fourier Transform of images represents elastic scattering events that (by definition) preserve energy and so in reciprocal space, they are located on the surface of the Ewald sphere and its Hermitian mate. In this precise description, the Fourier Transform of the complex value image needs to be placed on the Ewald sphere during the reconstruction process. For this reason, these complex, wavefront-corrected images are not just artificial constructs but rather they have fundamental meaning. In the traditional (back projection) method, the CTF function represents the approximation of reconstructing particles which are smaller than the depth of focus. This condition is resolution dependent and when the resolution becomes high enough that the depth of focus is smaller than the particle diameter, the CTF loses its meaning as a power spectrum modulation and instead, becomes a complex value function describing the phase of the scattered electron wavefront defining the image model and its inversion (3D reconstruction) (Russo and Henderson 2018).

Approximations used in cryoEM SPR are highly critical for computational efficiency, but they also affect the quality of the results. Here we presented the core aspects of novel data structures useful for computationally efficient SPR which is future-oriented, as it is compatible with more advanced computational layers which may be specific to particular approaches.

## Author Contributions

DB generated data; RB, YG, DB and ZO acquired experience with analyzing data; RB, YG, DB, and ZO wrote manuscript.

## Acknowledgments

We thank the Cryo-Electron Microscopy Facility (CEMF) at UT Southwestern Medical Center which is supported by grant RP170644 from the Cancer Prevention & Research Institute of Texas (CPRIT).

